# Examination of diurnal variation and sex differences in hippocampal neurophysiology and spatial memory

**DOI:** 10.1101/2022.03.12.484083

**Authors:** Lacy K. Goode, Allison R. Fusilier, Natalie Remiszewski, Jacob M. Reeves, Kavitha Abiraman, Matthew Defenderfer, Jodi R. Paul, Lori L. McMahon, Karen L. Gamble

**Affiliations:** Department of Psychiatry & Behavioral Neurobiology, University of Alabama at Birmingham Heersink School of Medicine, Birmingham, AL, USA; Research Computing, Information Technology, University of Alabama at Birmingham, Birmingham, AL, USA; Department of Cell, Developmental, & Integrative Biology, University of Alabama at Birmingham, Birmingham, AL, USA; Prescott Medical Communications Group, Chicago, IL, USA

## Abstract

Circadian rhythms are biological processes that cycle across 24 hours and regulate many facets of neurophysiology, including learning and memory. Circadian variation in performance on spatial memory tasks is well-documented; however, the effect of sex across circadian time remains unclear. Additionally, little is known regarding the impact of time-of-day on hippocampal neuronal physiology. Here, we investigated the influence of both sex and time-of-day on hippocampal neurophysiology and memory.

Performance on the object location memory (OLM) task depended on both circadian time and sex, with memory enhanced at night in males but during the day in females. Long-term synaptic potentiation (LTP) magnitude at CA3-CA1 synapses was greater at night compared to day in both sexes. Next, we measured spontaneous synaptic excitation and inhibition onto CA1 pyramidal neurons. Frequency and amplitude of inhibition was greater during the day compared to night, regardless of sex. Frequency and amplitude of excitation was larger in females, compared to males, independent of time-of-day, although both time-of-day and sex influenced presynaptic release probability. At night, CA1 pyramidal neurons showed enhanced excitability (action potential firing and/or baseline potential) that was dependent on synaptic excitation and inhibition, regardless of sex. This study emphasizes the importance of sex and time-of-day in hippocampal physiology, especially given that many neurological disorders impacting the hippocampus are linked to circadian disruption and present differently in men and women. Knowledge about how sex and circadian rhythms affect hippocampal physiology can improve the translational relevancy of therapeutics and inform the appropriate timing of existing treatments.

## Introduction

The hippocampus is the seat of learning and memory in the brain and its primary output is generated by the principal cells (i.e, pyramidal neurons) in area CA1. Action potential firing by a CA1 pyramidal neuron, like any other neuron, is a combined function of excitatory and inhibitory synaptic drive, intrinsic membrane properties regulating excitability, and neuromodulators (Spruston, 2008). A relatively unexplored facet in the hippocampus is how CA1 pyramidal neuron physiology is modulated by time-of-day. At the cellular level, time-of-day variations in biological function are generated by a transcriptional-translational feedback loop (Partch *et al*., 2014). Tissue-clocks throughout the body are hierarchically organized in a system that drives the timing of 24-h rhythms in physiology and behavior, enabling organisms to adapt to and anticipate regularly occurring events in their environment (Pilorz *et al*., 2020; Buijs *et al*., 2021). Circadian regulation of physiological processes is advantageous, and dysregulation of circadian rhythms can promote and exacerbate disease onset and symptoms (Logan & McClung, 2019; Colwell, 2021). Therefore, understanding circadian influence on physiology is crucial for designing interventions for diseases with circadian dysfunction, such as neurodegenerative diseases (Lee *et al*., 2021). Moreover, the majority of foundational knowledge concerning fundamental principles of hippocampal physiology is based on studies conducted in nocturnal, mostly male, rodents during their inactive phase (daytime). While the scientific community has begun to address the importance of sex as a factor in biomedical research, the importance of time-of-day is still relatively underemphasized. The overarching goal of this study was to begin to unveil how sex and time-of-day interact to influence daily variation in hippocampal physiology and function.

The suprachiasmatic nucleus (SCN) of the hypothalamus is the principal orchestrator of the endogenous circadian network, and electrical properties of SCN neurons vary across time-of-day. In fact, circadian regulation of neuronal excitability is widespread in the mammalian brain (Paul *et al*., 2020) and has been observed in a range of species, including rodents (Snider & Obrietan, 2018), Drosophila (Cao & Nitabach, 2008; Sheeba, 2008), and zebrafish (Elbaz *et al*., 2013). Although the SCN is the principal clock, autonomous circadian clocks exist in other brain regions, including hippocampus (Paul *et al*., 2020; Hartsock & Spencer, 2020). At the molecular level, subregions of hippocampus rhythmically express core clock proteins, with the cell body layer of area CA1 having the strongest expression of PER2 (Jilg *et al*., 2010). Moreover, over 600 genes, including those encoding ion channels and synaptic proteins exhibit circadian expression in hippocampus (Zhang *et al*., 2014; Renaud *et al*., 2015). At the cellular level, long term potentiation (LTP), a form of plasticity in which specific patterns of synaptic stimulation results in a long-lasting increase in the strength of synaptic transmission, is expressed at a greater magnitude at night compared to day in nocturnal mice (Chaudhury *et al*., 2005; Besing *et al*., 2017; Davis *et al*., 2020). Cognitive function is also regulated by the circadian system (Wright *et al*., 2012) and circadian regulation of performance on hippocampus-dependent memory assays has been demonstrated across several species (Snider *et al*., 2018). However, our understanding of how sex affects circadian regulation of cognition is limited. Furthermore, evidence at the cellular level is lacking, including a detailed understanding of how time-of-day and sex regulate synaptic drive onto and membrane properties of CA1 pyramidal cells. Here, we sought to determine how sex and time-of-day modulate the hippocampal circuit: from the behavioral level down to individual neuronal physiology. We found that circadian regulation of hippocampus-dependent memory is dependent on sex, while day-night differences in hippocampal LTP are not. We also found that synaptic transmission and neuronal excitability vary as a function of time-of-day and uncovered that some of these changes depend on sex.

## Methods

### Animals

All animal procedures followed the Guide for the Care and Use of Laboratory Animals, U.S. Public Health Service and were approved by the University of Alabama at Birmingham (UAB) Institutional Animal Care and Use Committee. All experiments used 6–12-week-old C57BL/6J mice of both sexes obtained from Jackson Laboratories (stock #000664) or from the C57BL/6J colony at UAB. Mice were maintained on a 12:12 light/dark cycle with ad libitum access to food (LabDiet Rodent 5001 by Purina) and water.

### Object Location Memory

The object location memory (OLM) task (Snider *et al*., 2016) was conducted under <10 lux dim red light. Four cohorts of mice were used, with each cohort consisting of eight males and eight females. Within each cohort, mice were assigned to undergo habituation, training, and testing either during the day or during the night. In cohorts 1 and 3, males were tested during the day, and females were tested at night. In cohorts 2 and 4, females were tested during the day and males were tested at night.

Mice were entrained to a 12:12 light/dark (LD) cycle, and then habituated to the arena for two days at either Zeitgeber Time (ZT) 4 or ZT16 (where ZT12 refers to lights off), for day and night, respectively (days 1-2, Fig. 1*A* and *B*). After day 2, mice were released into constant darkness (DD) and again habituated to the arena at projected circadian time (CT) 4 or 16 (days 3-4, where CT12 refers to the projected time of lights off from the prior LD cycle; Fig. 1*A* and *B*). Mice were allowed to acclimate to the behavior room for 20 min each day immediately prior to habituation or train/test. Habituation consisted of 5 min of handling followed by 5 min of arena exploration with visual cues present. Visual cues consisted of vertical stripes on one wall and a large red “X” on another wall. The arenas were 35.5 cm x 25.4 cm with 20.3 cm high walls. OLM training and testing occurred in DD, 24 h after the final day of habituation at projected CT 4 or CT 16. Objects were made with PRETEX™ Building Blocks (Item No.: 8030-100) and had three possible positions within the arena, all at least 8.9 cm away from the walls.

**Figure 1.**
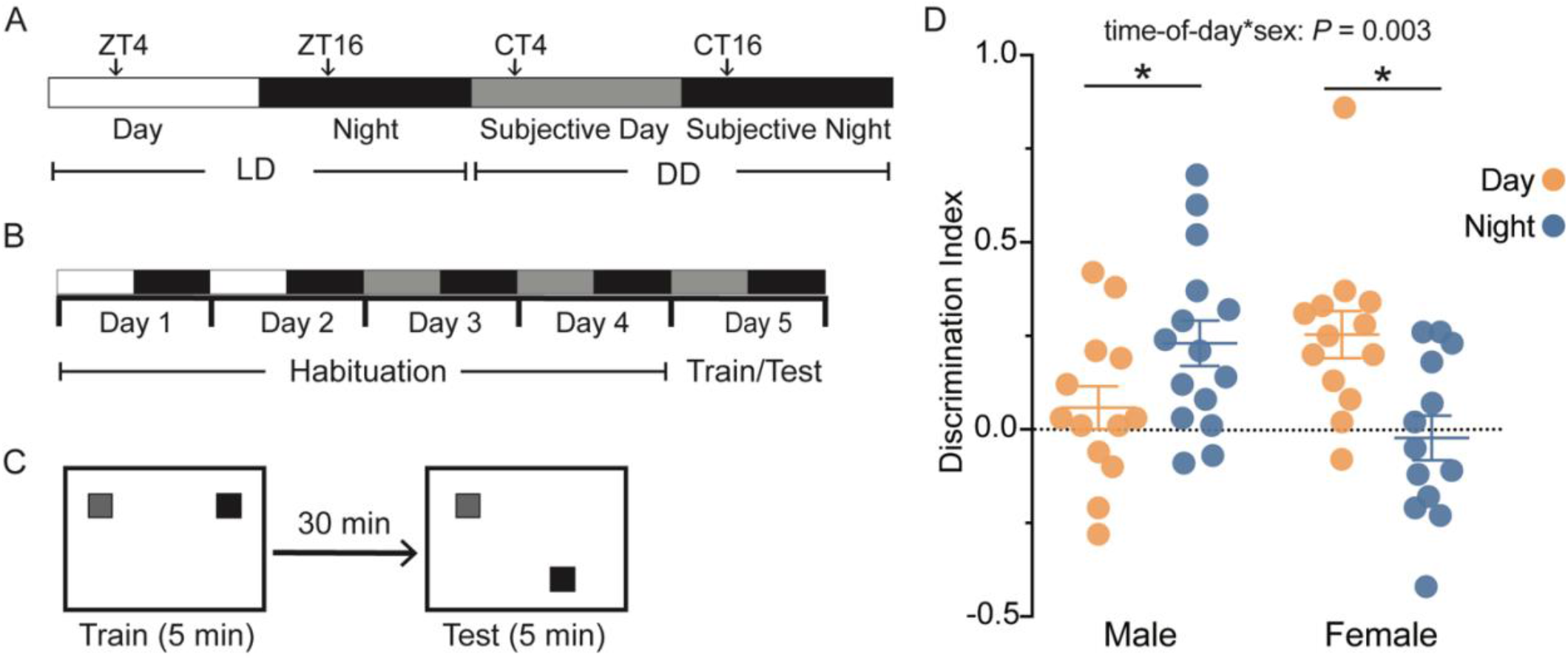
Circadian rhythms regulate object location memory (OLM) performance in a sex-dependent manner. *A*, schematic illustrating light-dark cycle (LD) and constant dark (DD) terminology. Zeitgeber Time (ZT) is used to denote time points in LD where ZT 12 refers to lights off. Circadian Time (CT) is used to denotate time points in DD where CT 12 is the projected night from the prior LD cycle. Experiments were conducted at ZT 4 or CT 4 (day) and ZT 16 or CT 16 (night). *B*, experimental timeline illustrating habitation during LD and DD on days 1-4 and training and testing procedures in DD on day 5. *C*, schematic of OLM experimental procedure. Mice were placed in arena and allowed to explore two objects during 5-min training phase at either CT 4 or CT 16. After a 30-min delay in the home cage, one object remained in the same location in the arena (familiar), and one object was moved to a new location (novel), and mice were placed back in the arena and allowed to explore for the 5-min testing phase. *D*, scatterplot displaying all individual Discrimination Index scores with mean ± SEM. Data were plotted for two time points and both sexes: male night, *n =* 13; male day, *n* = 13; female day, *n* = 15; female night, *n* = 13 (*interaction, *P* = 0.023, two-way ANOVA; *time-of-day for males, P = 0.047, simple main effects; *time-of-day for females, P = 0.003, simple main effects). In all plots, blue codes for observations at night, and orange codes observations made during the day. ZT, Zeitgeber Time; CT, Circadian Time; LD, Light-Dark; DD, constant darkness.

During training, each mouse was allowed to explore an arena with two objects for 5 min. Afterwards, the mouse was returned to its home cage for 30 min, during which one of the objects from the original exploration was moved to a new position (the novel location) while one remained in its original position (the familiar location; Fig. 1*C*). A 30-min recall period was chosen based on previously published methods (Snider *et al*., 2016) and to avoid memory interference due to sleep deprivation or memory enhancement from an overnight sleep period. In the test phase, each mouse was placed back in the arena with the novel and familiar location objects and allowed to explore for 5 min. All habituation, training, and testing were recorded at 30 FPS (ELP Camera Model: ELP-USBFHD05MT-KL36IR). Exploration was tracked using a computer model made via DeepLabCut. Object interaction was then analyzed using custom MatLab (MathWorks, Natick, MA) scripts developed by Dr. Mary Phillips (https://github.com/PhillipsML/DLC-NovelObject#dlc-novelobject). For data analysis, several exclusion criteria were applied: mice that exhibited a clear side preference, mice that spent most of their time exploring objects for the purpose of trying to escape the arena, and mice with a high preference to one object over the other during training were excluded. Discrimination index was calculated as: (time spent exploring novel object location—time spent exploring familiar object location)/(time spent exploring novel object location + time spent exploring familiar object location).

### Electrophysiology

#### Slice preparation

Mice were euthanized with cervical dislocation and rapid decapitation at ZT 0-1 or ZT 11-12 for day and night experiments, respectively. Both sex and time-of-day were interleaved. For extracellular field experiments, brains were removed and 350 µm coronal slices were prepared using a VT1200 S vibratome (Leica Biosystems) in an ice-cold solution containing the following (in mM): 85 NaCl, 2.5 KCl, 4 MgSO_4_ * 7 H_2_O, 0.5 CaCl_2_ * 2H_2_O, 1.25 NaH_2_PO_4_, 75 Sucrose, 25 NaHCO_3_, 25 Glucose saturated in 95% O_2_ and 5% CO_2_. Slices were allowed to rest for at least one hour in a recovery solution of standard artificial cerebral spinal fluid (ACSF) containing the following (in mM): 119 NaCl, 2.5 KCl, 1.3 MgSO_4_ * 7H_2_O, 2.5 CaCl_2_ * 2H_2_O, 1 NaH_2_PO_4_, 26 NaHCO_3_, and 11 glucose, bubbled with 95% O_2_/5% CO_2_. For whole-cell patch clamp experiments, brains were removed and 300-µm thick coronal slices were prepared using a VT1200 S vibratome (Leica Biosystems) in an ice-cold solution containing the following (in mM): 110 choline chloride, 25 glucose, 7 MgCl_2_, 2.5 KCl, 1.25 Na_2_PO_4_, 0.5 CaCl_2_, 1.3 Na-ascorbate, 3 Na-pyruvate, and 25 NaHCO_3_, bubbled with 95% O2/5% CO_2_. Slices were allowed to rest for at least one hour at room temperature in a recovery solution containing the following (in mM): 125 NaCl, 2.5 KCl, 1.25 Na2PO_4_, 2 CaCl_2_, 1 MgCl_2_, 25 NaHCO_3_, 25 glucose, bubbled with 95% O_2_/5% CO_2_. For experiments measuring inhibitory synaptic events, 2 mM kynurenic acid was added to the recovery solution.

#### Field recordings

Data were obtained from ZT 1-6 or ZT 12-18 for day and night recordings, respectively. Field excitatory postsynaptic potentials (fEPSPs) were recorded from slices in a submersion recording chamber continuously perfused with standard ACSF at 3-5 ml/min and 26–28°C. To stimulate Schaffer collateral axons, a bipolar stimulating electrode was placed in stratum radiatum within 200–300μm of a recording electrode. Data were acquired and analyzed using pCLAMP10/11(Molecular Devices, San Jose, CA). Data were recorded using a Kerr Scientific S2 amplifier (Kerr Tissue Recording System, Kerr Scientific Instruments, Auckland, NZ). Signals were digitized at 10 kHz (Digidata 1550B).

Input-output (I/O) curves were generated by measuring the slope of fEPSPs from CA1 stratum radiatum in response to a series of increasing stimulation intensities (0.2-200 µA, Δ10µA) at the Schaffer Collaterals. Baseline fEPSPs were obtained by delivering a 0.1Hz stimulation to elicit fEPSPs of approximately −0.20 mV/ms for 20 min.

Long term potentiation (LTP) experiments were conducted by obtaining and maintaining a stable baseline fEPSP response for 20 min, and then LTP was induced by delivering a high-frequency stimulation (HFS; 100Hz;0.5 s duration; delivered 2x with 15 s interval). This weaker stimulation protocol was chosen to avoid masking a day/night difference in LTP magnitude (Besing *et al*., 2017; Davis *et al*., 2020). After HFS, fEPSP slopes were recorded for 40 min.

#### Whole-cell patch clamp recordings

All data were collected from CA1 pyramidal neurons using the blind patch technique between ZT 2-6 (day) or ZT 13-17 (night) at 32°C in standard ACSF containing the following (in mM): 125 NaCl, 2.5 KCl, 1.25 Na_2_PO_4_, 2 CaCl_2_, 1 MgCl_2_, 25 NaHCO_3_, and 25 glucose, bubbled with 95% O2/5% CO_2_. Data were acquired using a Multiclamp 700B amplifier, Axon Digidata 1440A and 1550B digitizer, and pClamp10/11 software (Molecular Devices, San Jose, CA). Patch pipettes (BF150–086; Sutter Instruments, Novato, CA) were pulled on a Sutter P-97 horizontal puller (Sutter Instruments, Novato, CA) to a resistance between 2.5-5 MΩ. Cells were dialyzed for 5 min prior to experimental recordings. Cells used for analysis had access resistance < 30 MΩ that did not increase by >20% for the duration of each five-minute experiment.

For voltage clamp experiments, all cells were held at −70 mV and signals were filtered at 5 kHz and digitized at 10 kHz. Inhibitory postsynaptic currents (IPSCs) experiments used a patch pipette solution containing (in mM): 140 CsCl, 10 EGTA, 5 MgCl_2_, 2 Na-ATP, 0.3 Na-GTP, 10 HEPES, and 0.2% biocytin (pH 7.3, 290 mOsm), and 5 QX-314 (sodium channel antagonist) added at time of use. IPSCs were pharmacologically isolated with bath perfusion of 10μM NBQX (AMPAR antagonist, Hello Bio) and 5μM CPP (NDMAR antagonist, Hello Bio). Excitatory postsynaptic currents (EPSCs) experiments used a patch pipette solution containing (in mM): 100 CsOH, 100 Gluconic acid (50%), 0.6 EGTA, 5 MgCl_2_, 2 Na-ATP*3H_2_O, 0.3 Na-GTP, 40 HEPES, 7 Phosphocreatine, biocytin (0.2%), and 5 QX-314 added at time of use. EPSCs were pharmacologically isolated with bath perfusion of 10 µM Gabazine (GABA_A_R antagonist, Hello Bio). Separate experiments to measure miniature inhibitory and excitatory postsynaptic currents (mIPSCs/mEPSCs) were recorded as above with the addition 0.5 µM TTX (voltage-gated sodium channel inhibitor, Tocris).

For current clamp experiments, signals were filtered at 10 kHz and digitized at 20 kHz. Patch pipette solution contained (in mM): 135 K-Gluconate, 2 MgCl_2,_ 0.1 EGTA, 10 HEPES, 4 KCl, 2 Mg-ATP, 0.5 Na-GTP, 10 Phosphocreatine, and biocytin (0.2%) (pH 7.3, 310 mOsm, and 2–4 MΩ). Neuronal excitability was assessed by injecting progressive steps of depolarizing current from rest (0 pA to 500 pA at 20 pA increments) and counting the number of action potentials fired during each 1000ms current step. Baseline membrane potential was calculated as the mean voltage over the 1400 ms sweep during the 0pA step. Initial experiments were done in the absence of synaptic blockers to determine how sex and time-of-day contribute to CA1 pyramidal neuron excitability in the intact circuit. To determine how intrinsic excitability was affected by time-of-day and sex, a separate, follow-up experiment was conducted in the presence of 10μM NBQX, 5μM CPP, and 10μM Gabazine.

### Immunohistochemistry

#### Biocytin

To confirm that cells recorded to measure postsynaptic currents were CA1 pyramidal cells, all cells were filled with biocytin for at least 20 min. Slices containing filled cells were fixed in 4% PFA for at least 24 hours, then washed for 3 × 10 min in PBS, and incubated for 2-3 hours at RT in a TBS solution containing 10% NDS, 3% BSA, 1% Glycine, 0.4% Triton X, and streptavidin-488 (1:1000). Slices were then washed for 3 × 10 min in PBS and mounted on glass slides and coverslips with ProLong Gold Antifade mounting media containing DAPI. Slides were visualized on a BZ-X700 fluorescence microscope (Keyence). Any cells that could not be classified as CA1 pyramidal cells based on location and morphology were excluded from analysis.

#### Analysis and Statistics

Data were analyzed and visualized using SPSS (version 27/28) and Prism-GraphPad software. Assumptions of parametric tests, including normality and homogeneity of variance were assessed, and if violated, data were transformed, or nonparametric tests were used. Unless otherwise stated, significance was ascribed at *P* < 0.05.

#### Object location memory

OLM data were analyzed using an independent two-way Analysis of Variance (ANOVA) with time-of-day and sex as independent variables and discrimination index as the dependent variable.

#### Field recordings

Input-output data were analyzed using a linear mixed model with fEPSP slope as a function of Time-of-day, Sex, and Stimulation Intensity. For LTP experiments, the fEPSP slopes were normalized to the baseline responses, and responses obtained during the last 10 min of the 40-min post-HFS recording period were analyzed using a three-way ANOVA with repeated measures (RM-ANOVA).

#### Whole-cell patch clamp electrophysiology

Postsynaptic currents (inhibitory and excitatory) were automatically detected from a five-minute recording using pClamp’s event template search then manually inspected for false event detection. The amplitudes and interevent interval (IEI) were analyzed using a generalized estimating equation (GEE) that allowed parameter estimates with population-averaged models while taking into account correlations between repeated measures within subjects (Reed & Kaas, 2010; Cook *et al*., 2016). The GEE model specified an unstructured working correlation matrix structure, a subject effect of cell, and a within-subject effect of postsynaptic events. The raw data had a significant positive skew with extreme values and thus, were trimmed of upper and lower outliers (10%) followed by either a log transformation in the case of the amplitude data, or a log+1 transformation in the case of IEI data, in order to meet assumptions of normal distributions before analysis.

For current-clamp experiments, because data were collected from cells across the anterior-posterior axis of the hippocampus, and there are known differences in electrophysiological properties of CA1 pyramidal neurons across the dorsal-ventral axis (Spruston, 2008; Marcelin *et al*., 2012; Dougherty *et al*., 2012, 2013; Hönigsperger *et al*., 2015; Kim & Johnston, 2015; Malik *et al*., 2016; Milior *et al*., 2016), we divided cells into anterior (dorsal-like) or posterior (ventral-like) groups based on coronal section anatomy (Allen Reference Atlas from atlas.brain-map.org, Fig. 7A). When we included anterior-posterior axis as a factor in our initial ANOVA model, we found it contributed significantly to our results (p=0.005, main effect, four-way RM-ANOVA) and so we stratified the data accordingly. Action potentials were counted using the *Action Potential Counting* module in the Easy Electrophysiology software (Easy Electrophysiology Ltd, RRID:SCR_021190), with the default, *Auto-Threshold Spike* algorithm. Action potentials per current step were analyzed across current steps in which data did not violate assumptions of linearity and normality: 160pA-400pA. A three-way ANOVA with repeated measures revealed that sex was not a significant main effect in either the anterior or posterior dataset. As a result, sexes were pooled together and analyzed using a two-way ANOVA with repeated measures. Baseline membrane potential was calculated using pClamp, stratified based on the anterior-posterior axis, and analyzed using an independent two-way ANOVA.

## Results

### Day-night differences in OLM performance depend on sex

To examine the effect of sex on circadian rhythms of learning and memory, we utilized the object location memory (OLM) assay, which relies on a mouse’s tendency to explore objects in novel locations, to assess hippocampal spatial memory (Barker & Warburton, 2011; Takahashi *et al*., 2013; Chao *et al*., 2016). Though circadian and diurnal differences in performance on OLM have been reported (Takahashi *et al*., 2013; Snider *et al*., 2016), the effect of sex on diurnal variation in OLM performance remains poorly understood. We found that OLM performance varies across time-of-day; however, the pattern of diurnal difference in performance differed between sexes (*P =* 0.023, two-way ANOVA interaction). While males performed better at night compared to the day, as expected (*P =* 0.028, simple main effects comparing day vs night in males; Fig. 1*D*), female mice performed better during the day compared to night (*P =* 0.004, simple main effects comparing day vs night in females; Fig. 1*D*). There was no effect of time-of-day or sex on total exploration time (*P =* 0.926 and 0.936, two-way ANOVA main effects).

### LTP magnitude at night is greater than day, regardless of sex

Long-term potentiation (LTP) is considered a cellular correlate of learning and memory. LTP at CA3-CA1 synapses is higher at night compared to the day in male mice (Chaudhury *et al*., 2005; Besing *et al*., 2017; Davis *et al*., 2020), but to our knowledge, there are no published reports of the effects of time-of-day on LTP magnitude in adult female mice. Given our finding that diurnal differences in performance on a hippocampal-dependent memory assay are dependent on sex, we next sought to determine whether sex affects diurnal differences in LTP. First, to assess the strength of basal synaptic transmission at CA3-CA1 synapses, we generated I/O curves by measuring the fEPSP slope from CA1 stratum radiatum in response to Schaffer collateral stimulation over a series of increasing stimulation intensities (0.2-200 µA, Δ10µA) during the day and night in male and female mice (Fig. 2*A* and *B*). While neither sex nor time-of-day had a significant effect on basal synaptic transmission over the stimulus range tested (*P =* 0.552 and 0.981, respectively, LMM main effect), there was a significant sex by stimulation intensity interaction (*P* < 0.001, LMM; Fig. 2*A* and *B*). Males had larger fEPSP slopes compared to females only at 180µA,190 µA, and 200µA, regardless of time-of-day (*P =* 0.041, 0.043, and 0.035, respectively; simple main effects comparing males and females across all stim intensities; Fig. 2*A*). Overall, these results indicate that time-of-day does not affect basal synaptic transmission and that sex affects responses only at the highest stimulation intensities.

**Figure 2.**
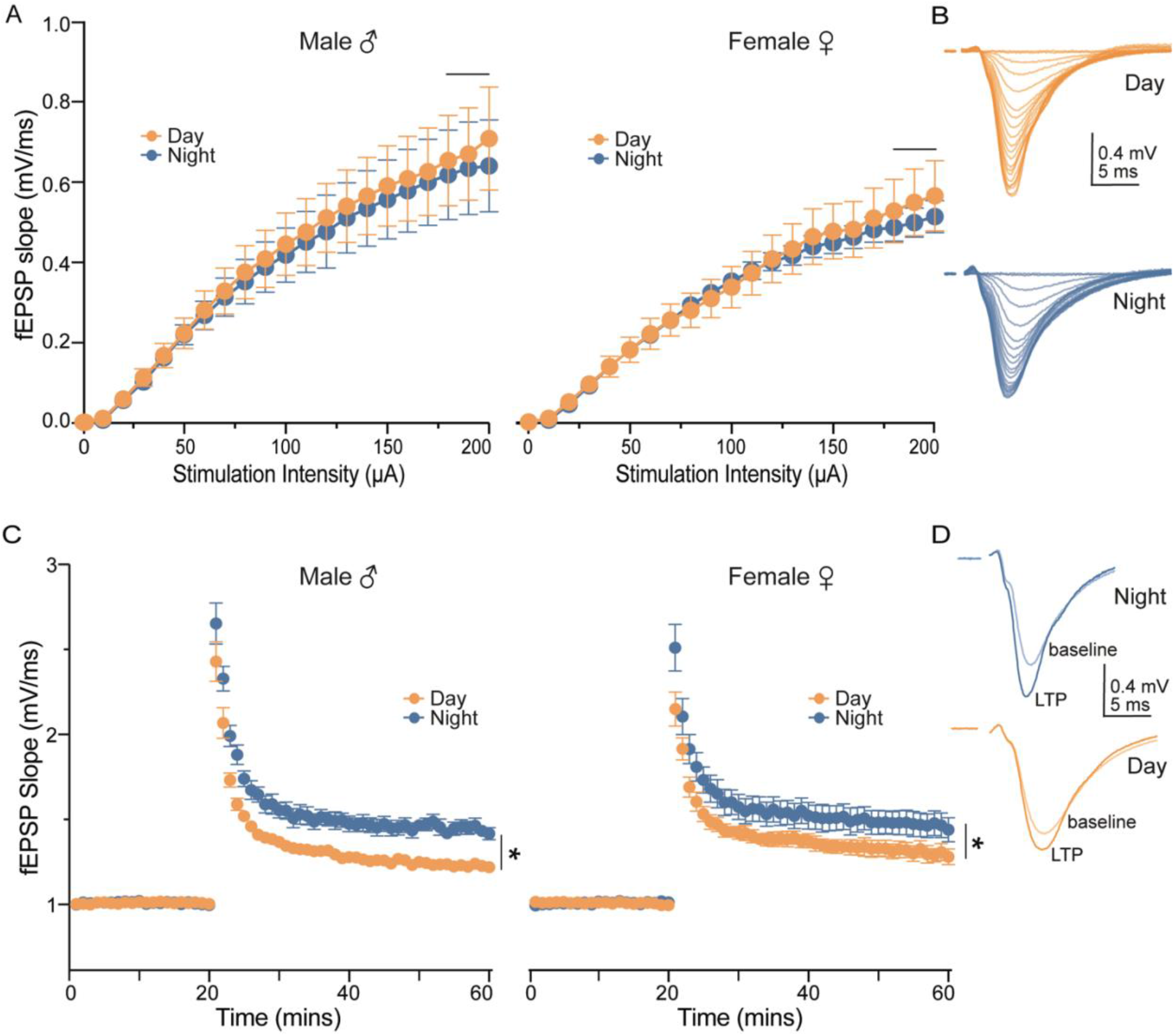
Long term potentiation (LTP) magnitude is greater at night compared to the day, regardless of sex. *A*, average slopes of fEPSPs in response to increasing stimulation of the Schaffer collaterals (time-of-day × sex interaction: *P* < 0.001). Note that significant sex differences in input-output responses were observed only at 180 µA (*P* = 0.041), 190 µA (*P* = 0.042), and 200 µA (*P* = 0.035), regardless of time-of-day, as indicated by horizontal line above stimulation intensities. Data were plotted for two time points and both sexes: male day, *n* = 13 slices from 3 mice; male night, *n* =10 slices from 3 mice; female day, *n* = 14 slices from 3 mice; female night, *n* = 12 slices from 3 mice. *B*, example fEPSPs from two male mice used to generate input-output curves in A. *C*, average slopes of fEPSPs before and after a HFS (100 Hz, 0.5s, 2X; t = 20 mins) to Schaffer collaterals (time-of-day: \**P* = 0.003; means ± SEM at 60 min, night: 1.451 ± 0.176; day: 1.286 ± 0.141). Data were plotted for two time points and both sexes: male day, *n* = 7 slices from 3 mice; male night, *n* = 8 slices from 3 mice; female day, *n* = 10 slices from 3 mice; female night, *n* = 9 slices from 3 mice. *D*, example fEPSPs from two female mice used to produce LTP in C. In all plots, blue codes recordings at night and orange codes recordings during the day. ♀ = female. ♂ = male. LTP, Long-term potentiation. fEPSPs, field excitatory postsynaptic potentials. HFS, high frequency tetanus. All statistical tests were performed with a three-way linear mixed model (input-output curves) or three-way, repeated measures ANOVA (LTP). Data are shown as means ± SEM.

Next, we assessed synaptic plasticity at the CA3-CA1 synapse by measuring LTP in response to a brief, high-frequency stimulation (HFS)(Fig.2*C* and *D*). As previously reported, the magnitude of LTP was greater at night compared to the day in both male and female mice (*P* = 0.003, three-way RM-ANOVA; Fig. 2*C* and *D*); however, there was no significant effect of or interaction with sex. Together, these findings suggest that time-of-day affects synaptic plasticity in male and female mice, without influencing basal synaptic strength.

### Synaptic inhibition onto CA1 pyramidal cells during the day is greater than night, regardless of sex

Changes in LTP can be attributed to synaptic mechanisms and/or intrinsic changes in excitability. Therefore, we first sought to determine if time-of-day and sex affect inhibitory and excitatory synaptic transmission onto CA1 pyramidal neurons. To examine synaptic inhibition onto CA1 pyramidal cells, we measured the amplitude and frequency of spontaneous inhibitory postsynaptic currents (sIPSCs) using whole-cell voltage clamp in male and female mice during the day and night (Fig. 3*A*). We found that sIPSC interevent interval (IEI) during the day was shorter than at night, regardless of sex (time-of-day: *P* = 0.033, GEE; Fig. 3*A* and *C*), indicating a greater frequency of inhibitory events during the day. The amplitude of sIPSCs during the day was larger than night in both males and females (time-of-day: *P* = 0.008, GEE; Fig. 3*A* and *E*). This increased day-time frequency and amplitude of sIPSCs suggest stronger inhibition onto CA1 pyramidal neurons during the day compared to night.

**Figure 3.**
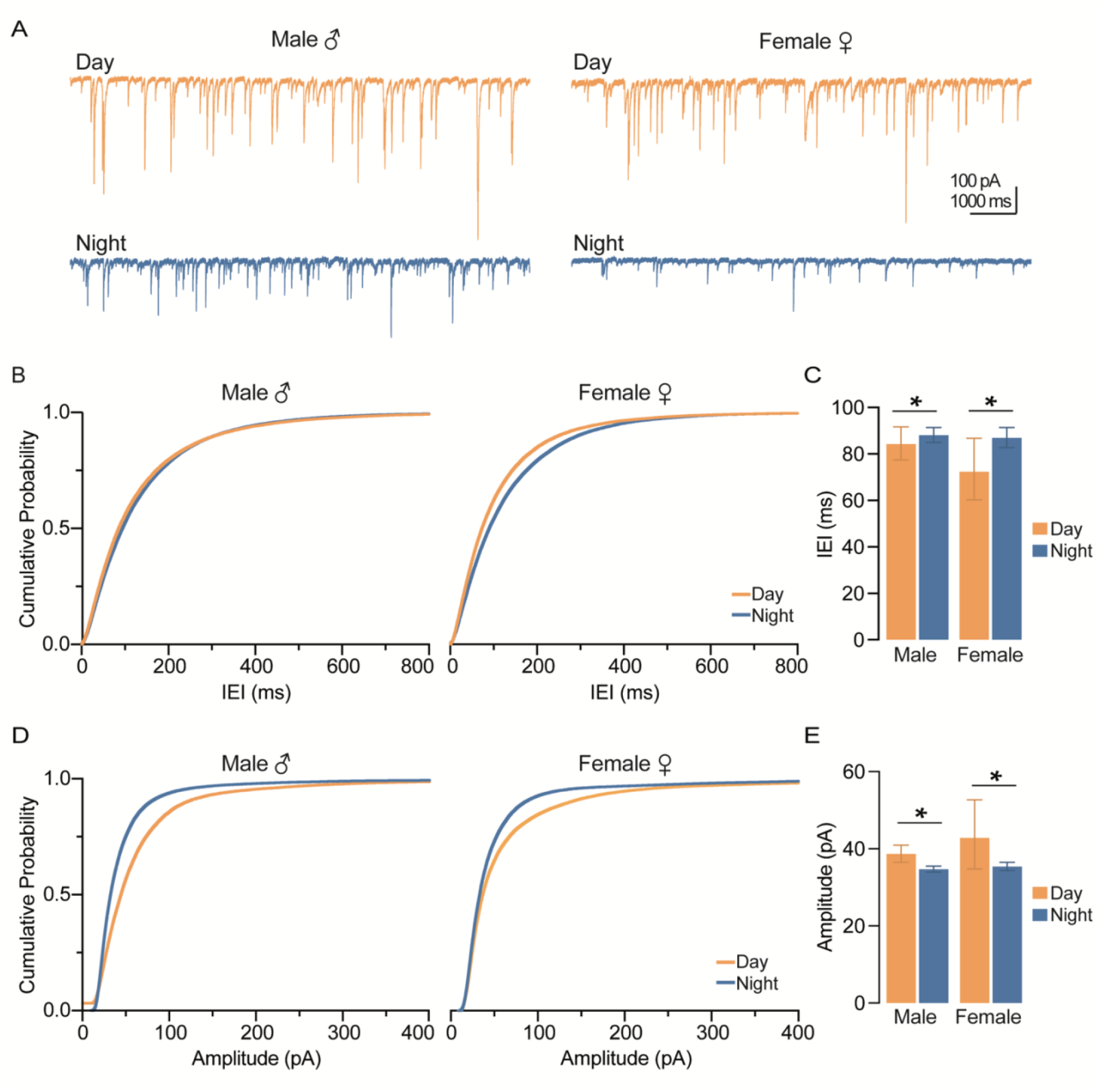
Action-potential-mediated inhibition onto CA1 pyramidal neurons is greater during the day compared to night, regardless of sex. *A*, example traces of IPSCs onto CA1 pyramidal neurons. Scale bars represent 100 pA and 1000 ms. *B*, cumulative probability distribution plots for the IEI of sIPSCs. C, estimated marginal means and confidence intervals of the IEI of sIPSCs (time-of-day: *P = 0.003; pooled EMM [95% confidence intervals] for night, 87.491 [82.215, 90.180] ms and day, 78.050 [72.164, 87.145] ms. *D*, cumulative probability distribution plots for the amplitude of sIPSCs. *E*, estimated marginal means and confidence intervals of the amplitude of sIPSCs (time-of-day: *P = 0.008; pooled EMM [95% confidence intervals] for night, 41.697 [37.523, 46.334] pA and day, 36.066 [35.424, 36.720] pA. sIPSCs were measured from 5 male mice during the day (*n* = 16 cells), 5 male mice at night (*n* = 28 cells), 5 female mice during the day (*n* = 18 cells), and 5 female mice at night (*n* = 17 cells). In all plots, blue codes recordings at night and orange codes recordings during the day. ♀ = female. ♂ = male. EMM, estimated marginal means. IEI, interevent interval. sIPSCs, spontaneous inhibitory postsynaptic currents. All statistical tests were performed with a two-way GEE model. Data were shown as EMM ± confidence intervals.

Stronger synaptic inhibition during the day could arise from an increase in presynaptic GABA release, or from increased postsynaptic GABA_A_R function. To distinguish between these possibilities, we measured miniature IPSCS (mIPSCs) in the presence of the voltage-gated sodium channel blocker tetrodotoxin (TTX) in both male and female mice during the day and night (Fig. 4*A*). While we found that neither sex (*P* = 0.392, GEE) nor time-of-day (*P* = 0.760, GEE) had a statistically significant effect on mIPSC IEI, events from males trended toward exhibiting a day-night difference (*P* = 0.068, sex*time-of-day interaction, GEE, Fig. 3C). The lack of day-night differences in females indicates that the time-of-day effects on sIPSCs is likely driven by local interneuron action potential firing. In males, the mean values between day and night differed by ∼12ms (mean and SEM: male day, 98.24 +/-1.05ms; male night, 85.78 +/- 1.04ms), suggesting that action-potential-independent inhibitory vesicle release may be more frequent at night (Fig. 3*C*). When we examined mIPSC amplitude, we unexpectedly found a significant interaction between time-of-day and sex (*P* = 0.038, GEE), with amplitudes in females being larger than males only during the day (*P* = 0.006, Wald Chi-Square pairwise comparisons, Fig. 4*E*); however, this ∼2pA difference is likely not biologically relevant (mean and SEM: female day, 34.68+/-1.01pA; male day, 32.67 +/- 1.02pA).

**Figure 4.**
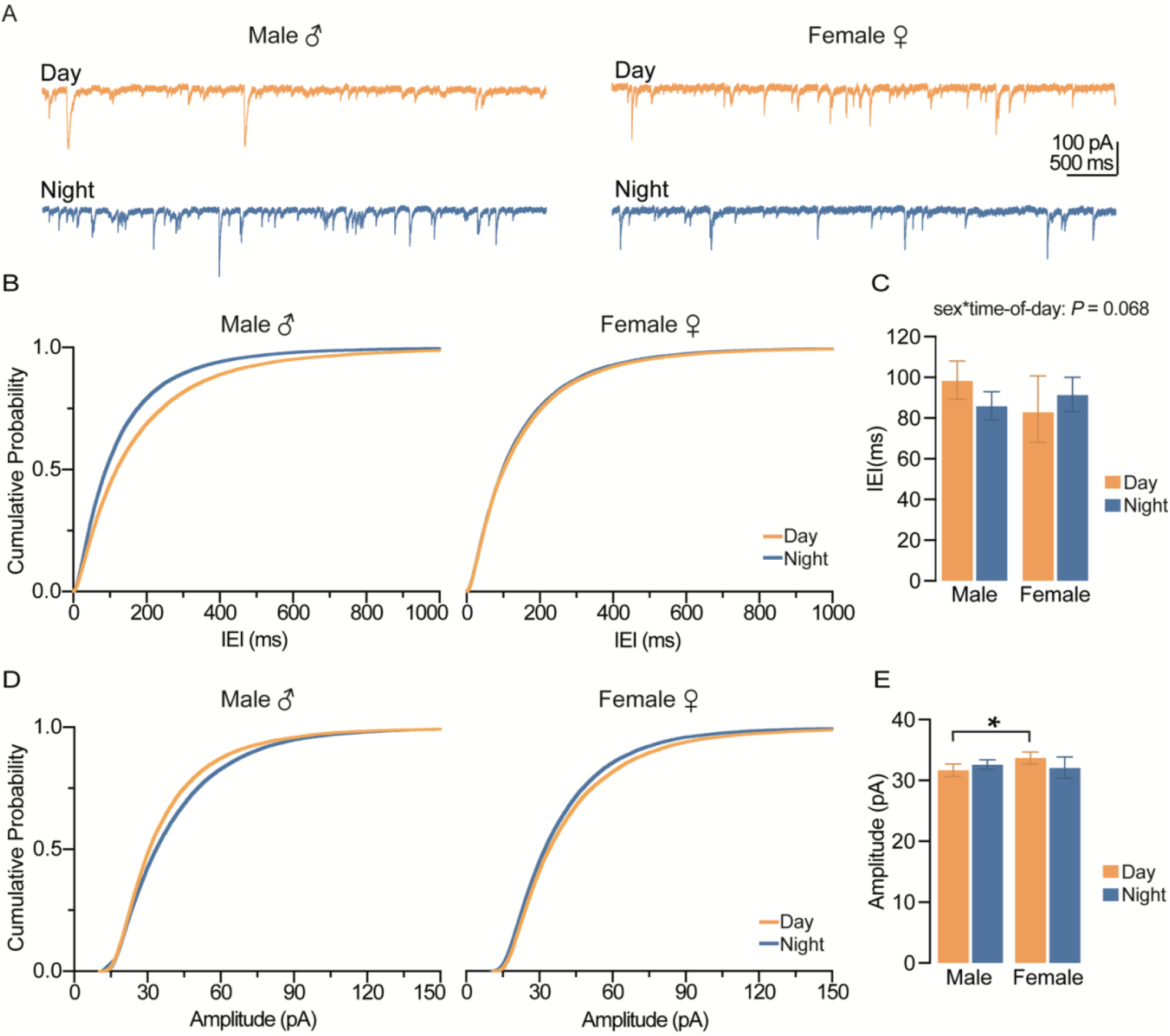
Action-potential-independent inhibition onto CA1 pyramidal neurons is influenced by sex and time-of-day. *A*, example traces of spontaneous, mIPSCs onto CA1 pyramidal neurons. Scale bars represent 100 pA and 500 ms. *B*, cumulative probability distribution plots for the IEI of mIPSCs. *C*, estimated marginal means and confidence intervals of the IEI of mIPSCs (interaction, *P* = 0.068). *D*, cumulative probability distribution plots for the amplitude of mIPSCs. *E*, estimated marginal means and confidence intervals of the amplitude of mIPSCs (interaction: *P = 0.038). Note that significant sex differences were observed during the day (*P* = 0.006) but not at night (*P* = 0.594). mIPSCs were measured from male 5 mice during the day (*n* = 12 cells), 5 male mice at night (*n* = 13 cells), 5 female mice during the day (*n* = 12 cells), and 5 female mice at night (*n* = 14 cells). In all plots, blue codes recordings at night and orange codes recordings during the day. ♀ = female. ♂ = male. EMM, estimated marginal means. IEI, interevent interval. mIPSCs, miniature inhibitory postsynaptic currents. All statistical tests were performed with a two-way GEE model followed by Wald Chi-Square pairwise comparisons. Data were shown as EMM ± confidence intervals.

Taken together, these spontaneous and miniature IPSC data suggest that action potential-dependent inhibition, but not spontaneous vesicle fusion, onto CA1 pyramidal cells is greater during the day compared to night in both males and females.

### Synaptic excitation onto CA1 pyramidal cells depends on sex

We next wanted to determine whether spontaneous excitatory synaptic input onto CA1 pyramidal neurons was affected by sex and time-of-day. First, we measured spontaneous excitatory postsynaptic currents (sEPSCs) using whole-cell voltage clamp recordings (Fig. 5*A*). Although there was no significant main effect of time-of-day on sEPSC amplitude (Fig. 5*E*), regardless of sex (*P* = 0.371, GEE), a statistical trend for a significant main effect of time-of-day indicated that sEPSC IEI recorded during the day may be greater than those recorded at night (*P* = 0.052, GEE), suggesting a greater frequency of excitatory events at night (Fig. 5*C*). Overall, we found that females had more excitatory synaptic input, with larger sEPSC amplitudes and shorter IEIs compared to males (*P* = 0.022 and 0.020, respectively, main effect of sex, GEE, Fig. 5*C,E*).

**Figure 5.**
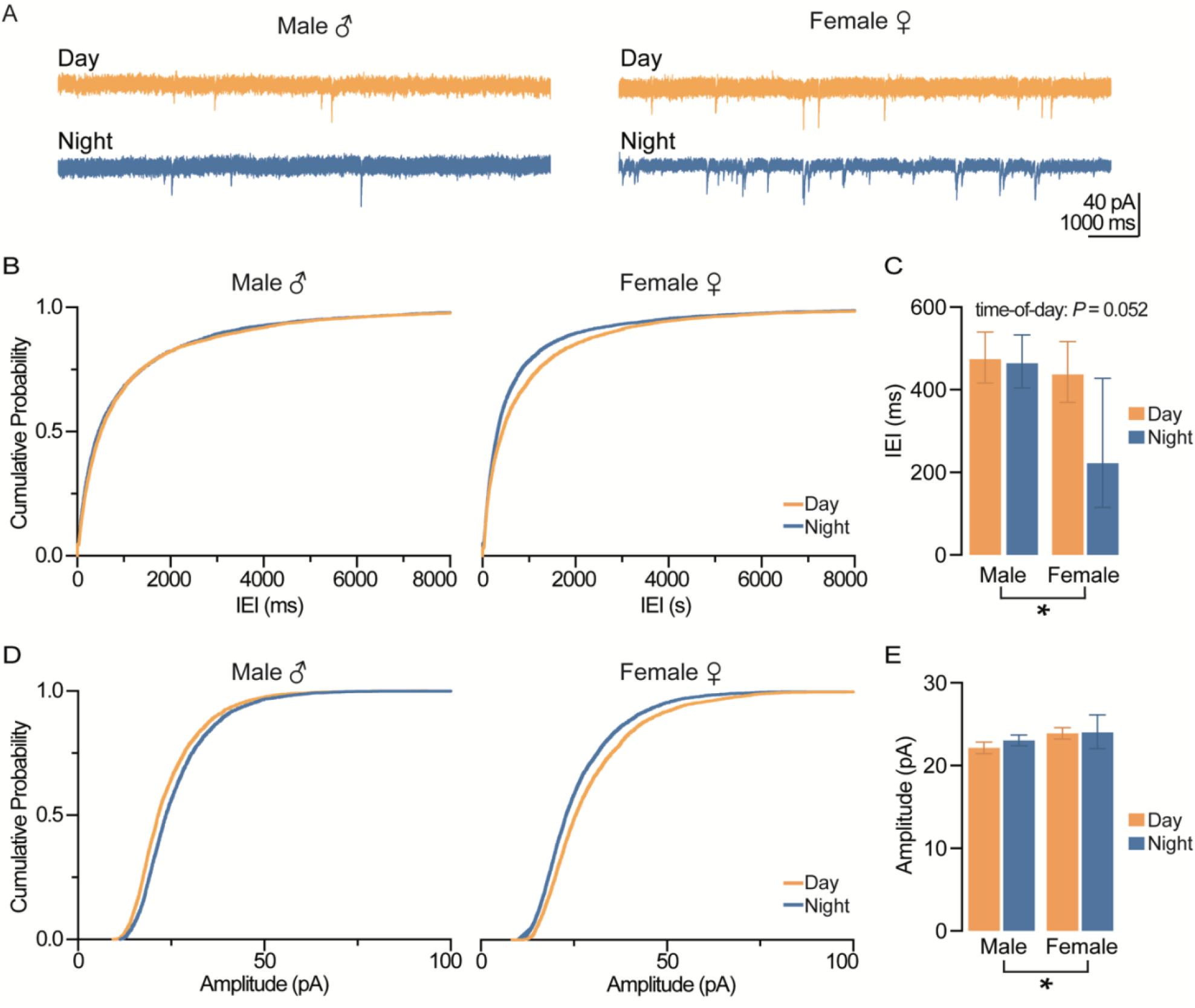
Action-potential-mediated excitation onto CA1 pyramidal neurons depends on sex. *A*, example traces of spontaneous sEPSCs onto CA1 pyramidal neurons. Scale bars represent 40 pA and 1000 ms. *B*, cumulative probability distribution plots for the IEI of sEPSCs. *C*, estimated marginal means ± confidence intervals of the IEI of sEPSCs (sex: \**P* = 0.022; pooled EMM [95% confidence intervals] for males, 469.002 [426.705, 515.530] ms and females, 311.536 [222.006, 436.968] ms. Note that differences in sEPSC IEIs recorded during the day (455.037 [410.344, 506.913]) and night (321.107 [230.600, 449.839] ms) failed to reach statistical significance (time-of-day: *P* = 0.052). *D*, cumulative probability distribution plots for the amplitude of sEPSCs. *E*, EMM and confidence intervals of the amplitude of sEPSCs (sex: \**P* = 0.020; pooled EMM [95% confidence intervals] for females, 24.952 [23.898, 26.046] pA and males, 23.578 [23.111, 24.058] pA. sEPSCs were recorded from 5 male mice during the day (*n* = 17 cells), 5 male mice at night (*n* = 16 cells), 5 female mice during the day (*n* = 16 cells), and 5 female mice at night (*n* = 18 cells). In all plots, blue codes recordings at night and orange codes recordings during the day. ♀ = female. ♂ = male. EMM, estimated marginal means. IEI, interevent interval. sEPSCs, spontaneous excitatory postsynaptic currents. All statistical tests were performed with a two-way GEE model. Data were shown as EMM ± confidence intervals.

Next, we repeated these experiments in the presence of the voltage-gated sodium channel blocker tetrodotoxin (TTX) and measured miniature spontaneous excitatory synaptic currents (mEPSCs) onto CA1 pyramidal neurons (Fig. 6*A*). Amplitude of mEPSCs did not vary across sex or time-of-day (*P* = 0.227 and *P* = 0.150, main effect of sex and time-of-day, respectively, GEE, Fig. 6*E*); however, day-night variation in mEPSC IEIs was dependent on sex (*P* = 0.021, time-of-day by sex interaction, GEE, Fig. 6*C*). In males, mEPSC IEIs were shorter at night than during the day (*P* = 0.002, Wald Chi-Square pairwise comparisons), indicating a greater frequency of excitatory events at night in male mice and therefore a likely increase in presynaptic release probability; however, there was no significant day-night difference in females (*P* = 0.765, Wald Chi-Square pairwise comparisons; Fig. 6*C*).

**Figure 6.**
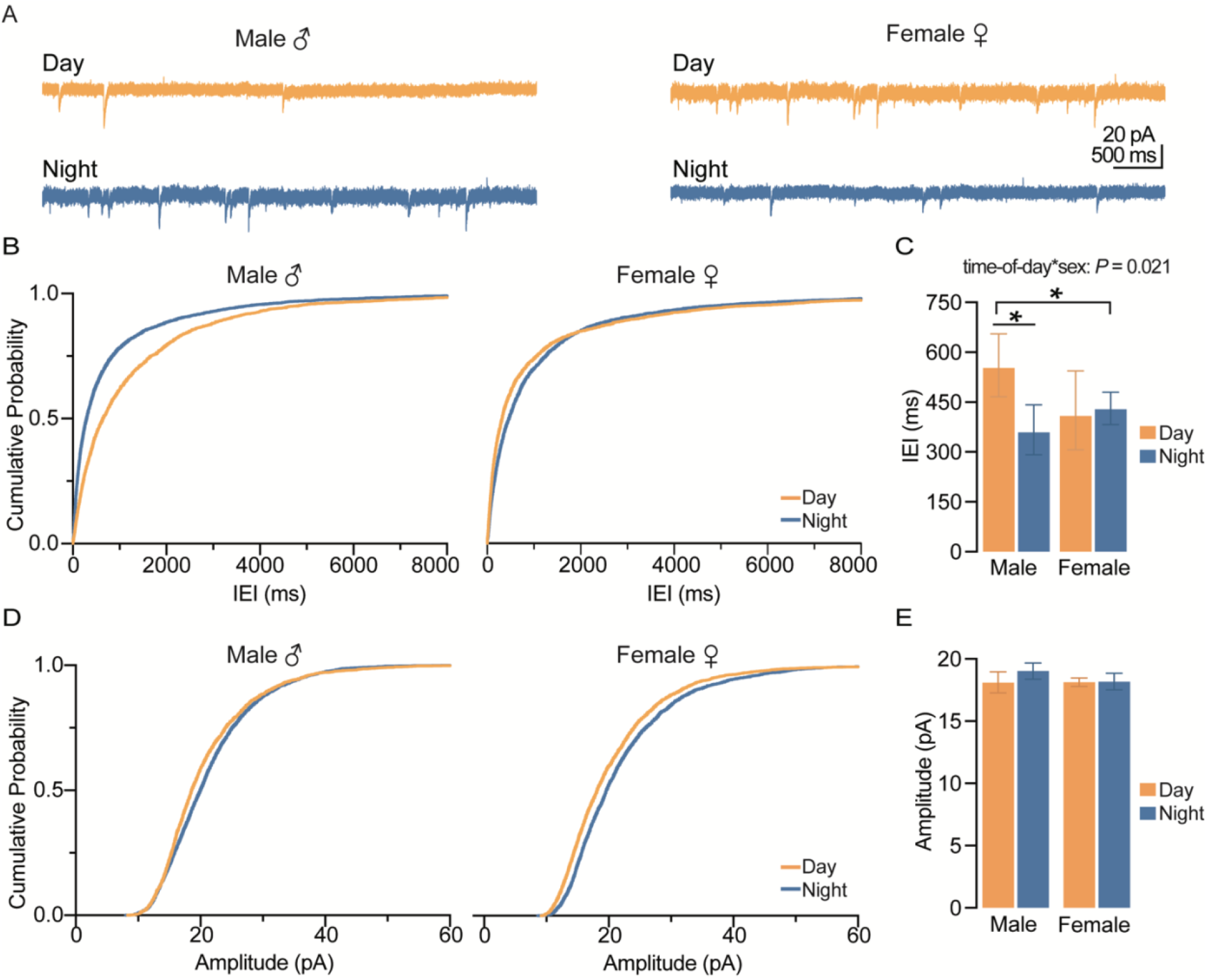
Action-potential-independent excitation onto CA1 pyramidal neurons is dependent on sex and time-of-day. *A*, example traces of spontaneous mEPSCs onto CA1 pyramidal neurons. Scale bars represent 20 pA and 500 ms. *B*, cumulative probability distribution plots for the IEI of mEPSCs. *C*, estimated marginal means and confidence intervals of the IEI of mEPSCs (interaction: \**P* = 0.021). Note that significant time-of-day differences were observed in males (\**P* = 0.002) but not in females (*P* = 0.765). *D*, cumulative probability distribution plots for the amplitude of mEPSCs. *E*, EMM and confidence intervals of the amplitude of mEPSCs (no significant effects). Spontaneous mEPSCs were recorded from 4 male mice during the day (*n* = 11 cells), 3 male mice at night (*n* = 13 cells), 3 female mice during the day (*n* = 14 cells), and 4 female mice at night (*n* = 13 cells). In all plots, blue codes recordings at night and orange codes recordings during the day. ♀ = female. ♂ = male. EMM, estimated marginal means. IEI, interevent interval. mEPSCs, miniature excitatory postsynaptic currents. All statistical tests were performed with a two-way GEE model followed by Wald Chi-Square pairwise comparisons. Data were shown as EMM ± confidence intervals.

**Figure 7.**
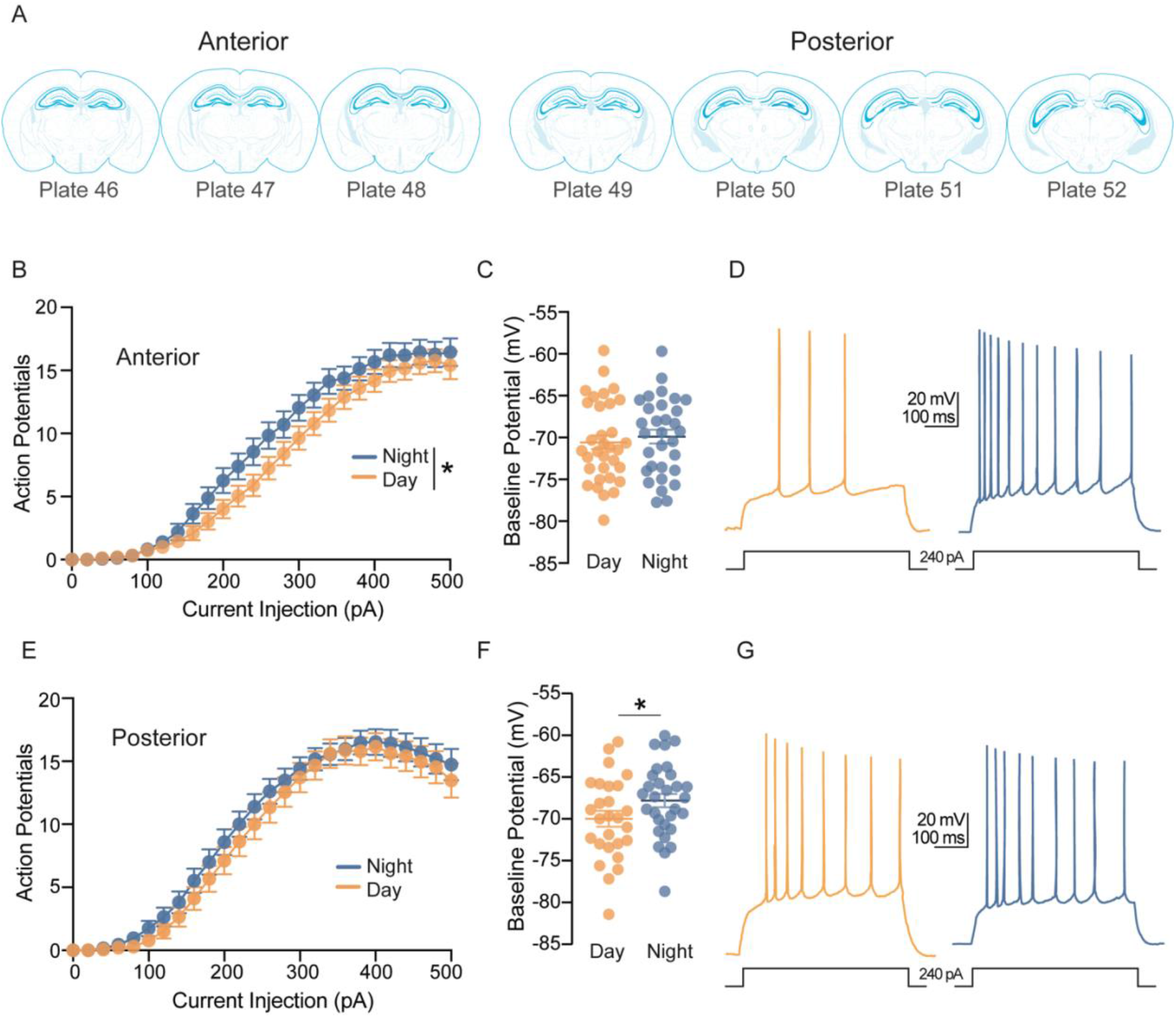
Excitability of CA1 pyramidal neurons depends on time-of-day but not sex. *A*, schematic locations of CA1 pyramidal neurons across the anterior-posterior axis in reference to the Allen Brain Atlas. Cells were considered “anterior” if recorded in hippocampal slices that corresponded to plate 48 or lower in the Allen brain atlas, and “posterior” if they corresponded to plate 49 or higher. *B*, average number of action potentials fired in response to increasing depolarizing current injections in neurons recorded from anterior slices from 14 mice at night (*n* = 33 cells) and 11 mice during the day (*n* = 33 cells; time-of-day: \**P* = 0.046). *C*, scatterplot of individual values with mean ± SEM of baseline membrane potentials of neurons recorded from anterior slices (time-of-day: ns *P* = 0.535). *D*, example traces of a 240-pA current step response in anterior slices from two male mice recorded during the day and night. Scale bars represent 20 mV and 100 ms. *E*, average number of action potentials fired in response to increasing depolarizing current injections in neurons recorded from posterior slices from 11 mice at night (*n* = 27 cells) and 11 mice during the day (*n* = 27 cells; time-of-day: ns *P* = 0.484). *F*, scatterplot of individual values with mean ± SEM of baseline membrane potentials of neurons recorded from posterior slices (time-of-day: \**P* = 0.044). *G*, example traces of a 240-pA current step response in posterior slices from two male mice recorded during the day and night. Scale bars represent 20 mV and 100 ms. In all plots, blue codes recordings at night and orange codes recordings during the day. All statistical tests were performed with a two-way repeated-measures ANOVA. Data were shown as means ± SEMs.

Taken together, these results suggest that the trend towards increased nighttime sEPSC frequency (especially in females) is action potential dependent. However, in the males, blocking action potentials uncovers a nighttime increase in frequency that was not seen in the sEPSCs.

### CA1 pyramidal neurons are more excitable at night

Broadly, our observations suggest that synaptic inhibition is greater at night and synaptic excitation is greater during the day; thus, we next wanted to determine if this opposing diurnal variation in synaptic excitatory and inhibitory input results in diurnal variation in CA1 pyramidal neuron excitability. To this end, we patched CA1 pyramidal cells in current clamp mode with the circuit intact (i.e., in the absence of synaptic antagonists) and without clamping cell membrane potential. We injected increasing amounts of depolarizing current (0 pA to 500 pA at 20 pA increments, 1000 ms duration) into pyramidal neurons and measured the number of action potentials elicited.

Because data were collected from cells across the anterior-posterior axis of the hippocampus, and there are known differences in electrophysiological properties of CA1 pyramidal neurons across the dorsal-ventral axis (Spruston, 2008; Marcelin *et al*., 2012; Dougherty *et al*., 2012, 2013; Hönigsperger *et al*., 2015; Kim & Johnston, 2015; Malik *et al*., 2016; Milior *et al*., 2016), we divided cells into anterior (dorsal-like) or posterior (ventral-like) groups based on coronal section anatomy (Allen Reference Atlas from atlas.brain-map.org, Fig. 7*A*). When we included the anterior-posterior axis as a factor in our initial ANOVA model, we found that it contributed significantly to our results (*P*=0.005, main effect, four-way RM-ANOVA) and we therefore stratified the data accordingly.

We found that sex had no significant effect on number of action potentials generated in response to depolarizing current injections of cells recorded from both anterior (*P* = 0.321, main effect of sex, three-way RM-ANOVA) and posterior slices (*P* = 0.568, main effect of sex, three-way RM-ANOVA), and thus, data from both sexes were pooled for final analysis. Anterior cells recorded at night fired more action potentials than those recoded during the day (*P* = 0.046, main effect of time-of-day, two-way RM-ANOVA, Fig. 7*B*). However, posterior cells displayed no statistical day-night difference in the number of action potentials fired (*P* = 0.484, main effect of time-of-day, two-way RM-ANOVA, Fig. 7*E*).

Examination of the baseline membrane potential revealed no differences in sex or time-of-day in cells recorded from anterior slices (*P* = 0.572 and 0.535, main effects of sex and time-of-day, respectively, two-way ANOVA, Fig. 7*C*). However, membrane potentials of cells recorded from posterior slices at night were more depolarized than those recorded during the day, regardless of sex (*P* = 0.044, main effect of time-of-day, two-way ANOVA, Fig. 7*F*).

Given that action potential firing and membrane potential are both measures of neuronal excitability, we can conclude that CA1 pyramidal neurons across the hippocampal circuit are more excitable at night, overall, but the mechanisms underlying this nighttime increase in excitability may vary across the hippocampal anterior-posterior axis.

### Day-night differences in CA1 pyramidal neuron excitability are not intrinsic

As the observed day-night variation in excitability could be driven by synaptic and/or intrinsic factors, we again assessed excitability in a separate cohort of animals in the presence of synaptic antagonists (CPP, NBQX, GBZ) to isolate the CA1 pyramidal neuron from the circuit. Under these conditions, we found no significant effect of time-of-day in the number of action potentials generated in response to depolarizing current injections in cells recorded from both anterior and posterior slices (*P* = 0.933 for anterior and *P* = 0.569 for posterior, main effect of time-of-day, two-way RM-ANOVA, Fig. 8*A* and *D*). Furthermore, baseline potentials under these conditions, were not affected by time-of-day or sex in cells recorded from either anterior (*P* = 0.896 and 0.888, main effect of sex and time-of-day, respectively, two-way ANOVA, Fig. 8*B*) or posterior slices (*P* = 0.999 and 0.702, main effect of sex and time-of-day, respectively, two-way ANOVA, Fig. 8*E*). Overall, these data suggest that increased nighttime excitability of CA1 pyramidal neurons is driven primarily by synaptic, not intrinsic, factors.

**Figure 8.**
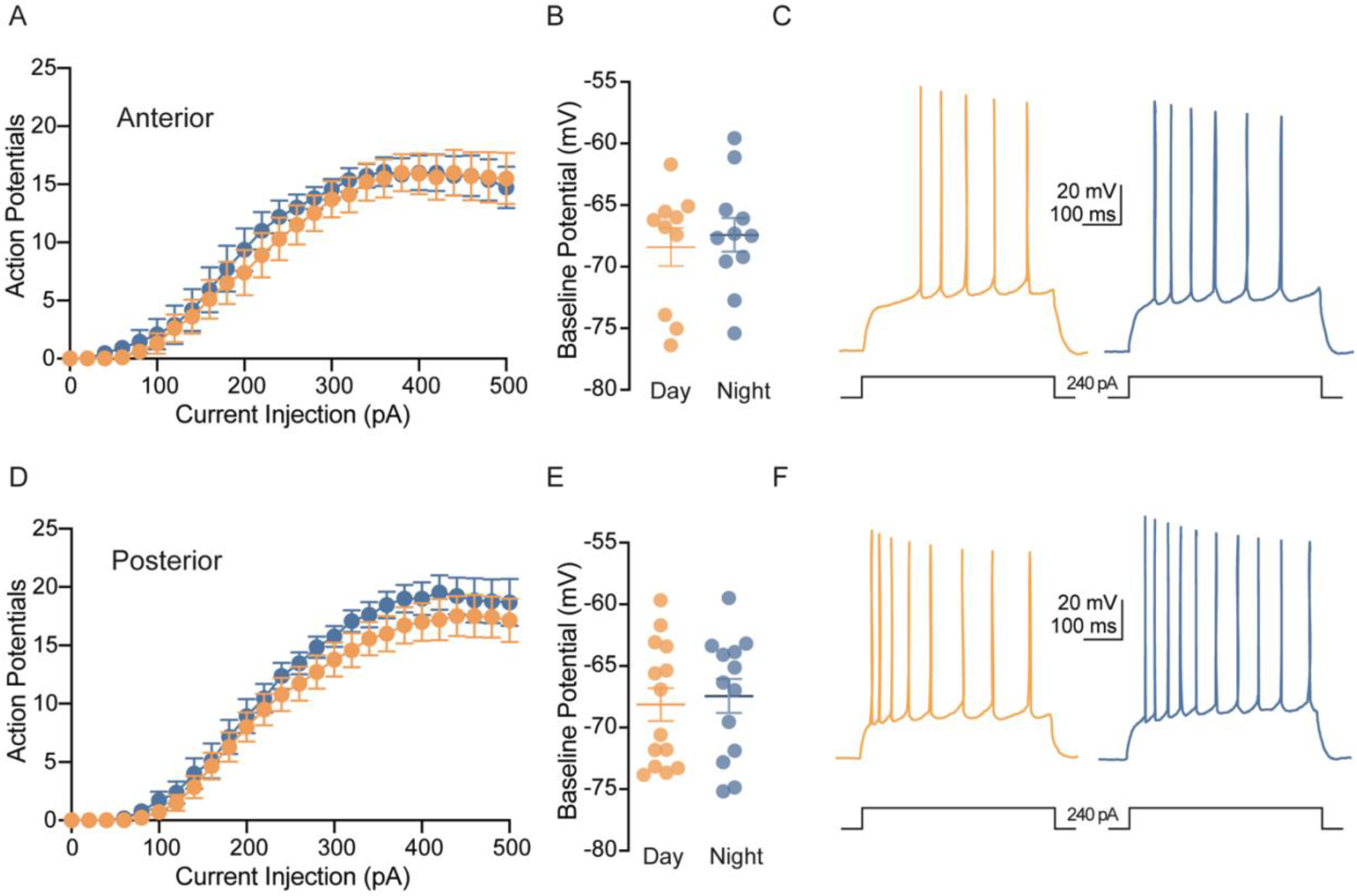
Diurnal differences in CA1 pyramidal neuron excitability are dependent on synaptic inputs. *A*, average number of action potentials fired in response to increasing depolarizing current injections in neurons recorded from anterior slices from 6 mice at night (*n* = 11 cells) and 6 mice during the day (*n* = 10 cells) in the presence of synaptic antagonists (time-of-day: ns *P* = 0.933). *B*, scatterplot of individual values with mean ± SEM of baseline membrane potentials of neurons recorded from anterior slices (time-of-day: ns *P* = 0.896). *C*, example traces of 240-pA current step response in anterior slices from two male mice recorded during the day and night in the presence of synaptic antagonists. Scale bars represent 20 mV and 100 ms. *D*, average number of action potentials fired in response to increasing depolarizing current injections in neurons recorded from posterior from 6 mice at night (*n* = 13 cells) and 6 mice during the day (*n* = 14 cells; time-of-day: ns *P* = 0.569). *E*, scatterplot of individual values with mean ± SEM of baseline membrane potentials of neurons recorded from posterior slices (time-of-day: ns P = 0.999). *F*, example traces of a 240-pA current step response in posterior slices from two male mice recorded during the day and night in the presence of synaptic antagonists. Scale bars represent 20 mV and 100 ms. In all plots, blue codes recordings at night and orange codes recordings during the day. All statistical tests were performed with a two-way repeated-measures ANOVA. Data were shown as means ± SEMs.

## Discussion

Here, we examined the effects of sex and time-of-day on multiple facets of hippocampal physiology, from behavior to individual neuronal physiology. We demonstrate that time-of-day impacts spatial learning and memory, LTP magnitude, synaptic inhibition onto CA1 pyramidal neurons, and CA1 pyramidal neuronal excitability. We found that sex was most influential on day-night differences in OLM performance, while its effect on synaptic transmission, LTP magnitude, and neuronal excitability were subtle or absent completely. Additionally, we found that position along the anterior-poster axis significantly impacts CA1 pyramidal neuron excitably. These findings illustrate the complexity of the hippocampal network and the importance of considering factors like sex and time-of-day in future studies.

While circadian rhythms regulate LTP and hippocampal-dependent learning and memory, the role of sex on diurnal differences in these processes was previously unknown. Surprisingly, we found that circadian rhythms regulate hippocampal-dependent memory performance in a sex-dependent manner. Male mice performed better on the OLM task at night, as previously reported (Takahashi *et al*., 2013; Snider *et al*., 2016). Female mice, however, performed better during the day. It is unclear why female mice would perform better during their inactive period but a possible explanation could be the estrous cycle, which was not controlled for in the present study. While more research in naturally cycling females is needed to make definitive conclusions about the specific impact of the estrous cycle on hippocampus-dependent spatial memory, studies in rats found that females in proestrus and estrous outperformed those in diestrus on object-recognition and object-placement tasks (Frye *et al*., 2007; Paris & Frye, 2008). Indeed, estrogen levels do impact performance on hippocampus-dependent memory tasks and administration of exogenous estradiol enhances performance on hippocampus-dependent memory tasks (Luine *et al*., 2003; Li *et al*., 2004; Phan *et al*., 2012; Vedder *et al*., 2013, 2014; Tuscher *et al*., 2015). Understanding how the estrous cycle and the circadian cycle converge to modulate cognition in females will be an interesting and important topic for future research.

Interestingly, we found that these sex effects on diurnal differences did not extend to diurnal regulation of LTP, which is considered a cellular correlate of learning and memory. This could be an example of how the same physiological process (LTP) is used to achieve different outcomes, depending on context (males vs females). Additionally, while the OLM task we chose is hippocampus-dependent (Barker & Warburton, 2011), learning and memory are complex processes that can rely on multiple memory systems, and perhaps females rely more heavily on other circuits compared to males. Moreover, the estrous cycle can influence learning strategies and the relative contributions of different memory circuits in females (Korol *et al*., 2004). Therefore, it is entirely possible that the sex-dependent effects we observed in the OLM task are mediated by a mechanism other than LTP at the CA3-CA1 synapse. Another factor that could explain our findings is our use of a high frequency stimulation, as opposed to a more physiological stimulation (i.e., theta-burst), which may have occluded detection of sex-dependent regulation of LTP. Regardless, these results exemplify the importance of accounting for both sex and time-of-day when designing research studies and interpreting results.

Previous work from our laboratory revealed that sIPSCs onto CA1 pyramidal neurons exhibit diurnal differences and that those diurnal differences are lost in a mouse model of Alzheimer’s Disease (Fusilier *et al*., 2021). Here, we replicated and expanded on previous findings by examining sIPSCs in both sexes and conducting additional experiments in the presence of TTX (mIPSCs) to begin to identify pre- and post-synaptic mechanisms contributing to diurnal differences. The lack of diurnal differences in mIPSCs IEI suggest that that action potential-dependent inhibition onto CA1 pyramidal cells is greater during the day than night in both male and female mice. Given that these inhibitory currents were pharmacologically isolated with glutamate receptor antagonists, it is likely that increased daytime interneuron activity is spontaneously generated. Indeed, prior reports suggest that some interneurons in area CA1 are spontaneously active (Sik *et al*., 1995; Maccaferri & McBain, 1996; Oliva *et al*., 2000; Amilhon *et al*., 2015; Huh *et al*., 2016; Miri *et al*., 2018); however, definitive evidence of diurnal variation in spontaneous interneuron firing in hippocampus is lacking and will be an important area of future study that could provide insight into circadian dysfunction associated with diseases involving hyperexcitability of the hippocampal network, including Alzheimer’s disease and epilepsy.

In addition to receiving inhibitory input from local interneuron populations, major excitatory input onto CA1 pyramidal cells arrives from the axons of principal neurons of the downstream area CA3 (Schaffer collaterals; CA3 pyramidal cells) or from the entorhinal cortex (temperoammonic pathway). Examination of sEPSCs revealed a trend for excitatory input onto CA1 pyramidal cells to be greater at night than during the day. In the absence of action-potential dependent neurotransmitter release, this phenomenon persists in males, but not in females, suggesting that increased nighttime excitation in females is likely action-potential driven. CA3 pyramidal cells also exhibit diurnal differences in excitability, such that night cells exhibit larger calcium current, decreased afterhyperpolarization, and reduced spike frequency adaptation compared to day cells (Kole *et al*., 2001). This increased nighttime CA3 pyramidal cell excitability could translate to increased sEPSC onto CA1 pyramidal cells at night compared to day.

We next wanted to examine how diurnal variation in excitatory and inhibitory synaptic transmission might impact CA1 pyramidal neuron excitability. A previous study in rats found that membrane excitability oscillated across circadian time, with neurons being more depolarized during the subjective late night/subjective early day (Naseri Kouzehgarani *et al*., 2020). We recently reported increased nighttime excitability in mouse CA1 pyramidal neuron excitability (Fusilier *et al*., 2021). Here, we replicated our previous findings and expanded our study to account for potential sex differences and the influence of the anterior-posterior hippocampal axis. Overall, we found that, in an intact synaptic circuit (i.e., without synaptic antagonists), CA1 pyramidal neurons were more excitable at night compared to day, regardless of sex. Interestingly, nighttime enhancement of excitability was not uniform across the hippocampal anterior-posterior axis. While neurons recorded from anterior slices fired more action potentials in response to depolarizing current injections and displayed no diurnal difference in baseline membrane potential, posterior neurons were more depolarized at night but did not display a statistically significant nighttime increase in number of action potentials fired. These findings suggest that the underlying mechanisms for nighttime enhancement of neuronal excitability may be different depending on location across the anterior-posterior axis. While our coronal slice preparation meant we were unable to truly isolate ventral hippocampus from dorsal hippocampus, we observed that neurons from posterior slices displayed characteristics consistent with previously published data collected from ventral CA1 pyramidal neurons, while neurons from anterior slices were consistent with data examining dorsal CA1 pyramidal neurons (Malik *et al*., 2016; Milior *et al*., 2016). Specifically, posterior neurons were more excitable compared to anterior neurons, reaching max firing rate earlier than anterior neurons. Given known differences in dendritic morphology and ion channel expression in dorsal vs ventral hippocampal neurons (Bannerman *et al*., 2004; Fanselow & Dong, 2010; Marcelin *et al*., 2012; Dougherty *et al*., 2012, 2013; Hönigsperger *et al*., 2015; Kim & Johnston, 2015; Malik *et al*., 2016; Milior *et al*., 2016; Soltesz & Losonczy, 2018), it is unsurprising that mechanisms underlying diurnal differences in excitability may be different across these populations. The absence of day-night differences of neuronal excitability in the presence of synaptic antagonists indicates that this effect is driven mostly by synaptic, as opposed to intrinsic, properties. Additionally, the difference between anterior and posterior cells was lessened in the presence of synaptic antagonists, suggesting synaptic factors could be at least partially responsible for some of the regional differences we observed. It will be interesting to narrow down how the expression and function of various neurotransmitter receptors and ion channels are modulated across both time-of-day and location along the longitudinal axis.

The hippocampus is one of the most studied and well-characterized circuits in the mammalian brain. However, most of the knowledge about how this circuit functions is based on studies conducted during the day in, mostly male, nocturnal rodents. Failing to account for factors like time-of-day and sex, leads to an incomplete picture hippocampal physiology and how it dynamically functions across multiple contexts. Sex and time-of-day are especially important considerations for the translational relevancy of studying the hippocampus in models of diseases like Alzheimer’s or epilepsy, which are influenced by circadian rhythms and can affect men and women differently. It is important to note that, except for experiments testing OLM, all experiments in this study were conducted in a light-dark cycle. Therefore, future studies in constant conditions are needed in order to determine the role of the circadian system on observed diurnal differences. While only two timepoints were examined here, it will be interesting and helpful to determine how hippocampal physiology is dynamic across multiple time points in the circadian cycle in future studies. In conclusion, this study reveals diurnal variation in hippocampal synaptic and neuronal function, and underscores the importance of considering sex, circadian rhythms, and neuronal heterogeneity within a brain region in the study of neural circuits.

## Acknowledgments

The authors would like to thank Animal Resources Program at UAB, Camille Smith, and Ananya Swaroop for their support in managing, maintaining, and caring for the animals used in this study. The authors gratefully acknowledge the resources provided by the University of Alabama at Birmingham IT-Research Computing group for high performance computing (HPC) support and CPU time on the Cheaha compute cluster.

## Author Contributions

All experiments were conducted at the University of Alabama Birmingham. AF, LG, NR, and JR performed experiments and analyzed data. MD and KG assisted with data computation and statistical analysis. LM participated in the discussion and interpretation of results. AF and LG wrote the article; KG, KA, JP, and LM revised it. All authors have approved the final version of the article and agree to be accountable for all aspects of the work. All persons designated as authors qualify for authorship, and all those who qualify for authorship are listed. Note: KA’s contribution to this work was independent from their role with Prescott Medical Communications Group.

